# Bacteriophage treatment of *Pseudomonas aeruginosa* PA14 infection does not alter the native microbiome of *Caenorhabditis elegans*

**DOI:** 10.64898/2026.07.16.738890

**Authors:** Jennifer E. Dumaine, Aditya Mantha, Alexander Ngo, Andrea Choi, Eric Song, Similoluwa Olaniyi, Aidan Tran, Katherine Herbert, Aaron Quijije, Haley Shaw, Allison Gash, Shreyan Mitra, Andrew Ho, Chris Kovacs, Car Reen Kok, Nicholas Be, F. John Burpo, Andrew R. Kick

## Abstract

Bacteriophage therapeutics are alternatives to antibiotics for treating multi-drug-resistant (MDR) bacterial infections. Bacteriophages kill host bacteria species with targeted precision, unlike the broad-spectrum nature of antibiotics. While the precision and the natural occurrence of phages in the environment have drawn attention to phages as new “drugs” in the time of accelerated antibiotic resistance, it remains poorly understood how phage treatment impacts the community of organisms composing the microbiome. Before phage therapies can be more widespread in practice, this understanding is required. *Caenorhabditis elegans* lends itself as an excellent model organism to address this question, as worm populations are raised under sterile conditions and the gut microbiome can be populated experimentally by the worms’ natural bacterivore diet. Here we have established a *C. elegans* infection and phage treatment model using fluorescent *Pseudomonas aeruginosa* PA14 to quantify infection. We isolated and whole genome sequenced PA14-specific lytic phage from wastewater samples near West Point, New York. To characterize the impact of treatment on the microbiome, we populated the *C. elegans* gut microbiome with 11 strains of the native *C. elegans* microbiome (CeMbio) and used PacBio sequencing of the full length 16S rRNA gene to characterize the microbiome during PA14 infection in the presence and absence of phage treatment (*n* = 100 worms per sample, 4 replicates per condition). Taxonomic classification, phylogenetic analysis, and diversity analysis performed using a HiFi 16S bioinformatics pipeline revealed that phage treatment and infection level did not change the overall composition of the *C. elegans* gut microbiome.

**Importance:** Multidrug resistant infections are emerging faster than new antibiotic compounds can be synthesized. Bacteriophages are natural viruses of bacteria that seek out and kill only their host bacteria species. Bacteriophage therapeutics are a promising solution for drug resistant infections in patients where no other treatment exists. However, while phage treatment is effective, there is little understanding about what happens within the complex community of the microbiome when a phage treatment is employed. Here we establish a new model utilizing *C. elegans* to integrate a multidrug resistant infection, phage treatment, and the native microbiome of the *C. elegans* worm to address how the microbiome changes in response to phage treatment. We find that the microbiome remains unchanged after bacteriophage treatment, highlighting the targeted specificity of phage therapy and suggests fewer side effects for phage therapy than antibiotic treatment for the same infection.

## Introduction

Multidrug Resistant (MDR) infections are recognized by the World Health Organization (WHO) as an emergent global health concern [1]. There has been a 43% increase in MDR infections around the world since 2025, with the largest increase occurring in healthcare-associated infections (67% increase) [2]. Widespread drug resistance is increasing faster than the discovery of new antimicrobial compounds, accelerating the problem surrounding resistance [1, 3, 4]. Bacteriophage therapeutics present a promising alternative to antibiotic use, as the targeted specificity of phages and their natural occurrence both in the environment and the microbiome make these attractive candidates for use. Bacteriophage therapeutics are not a new concept, but historically the advent of antibiotics replaced phage therapy in the Western World [5]. As antimicrobial resistance continues to rise, phage therapy is an innovative solution for this worsening problem.

Phages are highly specific viruses that infect and kill bacteria. Lytic phages infect, replicate, and lyse host bacteria, leaving other cells unharmed. Phages have been shown to resensitize antibiotic-resistant bacteria to drug treatment, providing another solution to the ongoing MDR problem [3, 6, 7]. Phage therapy begins with identifying phages targeting the pathogen of interest, which may be difficult for novel infections or multi-organism infections. Bacteria may develop phage resistance, requiring the use of phage cocktails for complex infections [3, 8]. More recently, synthetic phages have been engineered to overcome host range limitations, to enhance lytic activity and combat antibiotic resistance; however, it is a laborious process and there is also limited specificity, a risk to develop resistance and regulatory and clinical hurdles to overcome [9, 10]. The targeted precision of phage therapy is particularly attractive in the fight against ESKAPE pathogens (*Enterococcus faecium, Staphylococcus aureus, Klebsiella pneumoniae, Acinetobacter baumannii, Pseudomonas aeruginosa*, and *Enterobacter species*), which are most associated with recalcitrant bacterial infections and are currently facing diminishing options for treatment [11].

While phage therapies are effective in eliminating their target host species, little is known about the effects of phage treatment on the host microbiome. The community of organisms composing the microbiome provides direct competition for space and resources as part of the innate immune response [12]. Colonization and later resolution of pathogen infection may cause changes in the overall microbiome community structure. Treatment with broad spectrum antibiotics not only targets pathogens, but also kills “good” bacteria of the microbiome, allowing for outgrowths of fungus or opportunistic infections such as *Clostridium difficile* [4]. In contrast, phage therapies target only their host organism, though removing one organism from a complex ecological community may have unintended consequences due to shifts in available resources [12, 13]. The impact of phage administration on the microbiome has been explored in gnotobiotic mouse models, but these studies are limited by reconstituting the mouse gut with representative members of the human microbiome and not the entire community [14, 15]. Here we establish a new model to investigate the impact of phage therapy on the gut microbiome when used to treat an MDR infection in *Caenorhabditis elegans,* such that the host, pathogen, microbiome, and phage therapeutic are all within the same organism*. C. elegans* is an ideal model organism as it is maintained under germ free conditions, the bacterivore diet provides control over gut colonization, and the complete worm gut microbiome has been identified and is available for purchase [16]. *Pseudomonas aeruginosa* PA14 is an ESKAPE pathogen that has been well characterized within the *C. elegans* model [17, 18]. Here we use a fluorescent protein expressing PA14 strain and PA14-specific bacteriophage isolated from wastewater samples near West Point, NY, to investigate phage-bacteria interactions within the same host. We colonized the gut microbiome *of C. elegans* using the commercially available worm microbiome (CeMbio) and uncovered that the presence of the microbiome alone reduces PA14 infection burden. We next used 16S sequencing technology to analyze changes in the microbiome composition and found that bacteriophage treatment did not significantly alter the microbiome, regardless of treatment status or infection level. Overall, we demonstrate the targeted specificity of phage therapy with limited consequence to the existing microbiome.

## Results

### Generation of a Fluorescent PA14 infection model in *C. elegans*

*Caenorhabditis elegans* is a well-established model for studying gut innate immunity [19,20]. Bleach synchronization sterilizes the gut, which can then be colonized with defined bacterial communities through feeding [16,21]. *Pseudomonas aeruginosa* PA14 has been a model for pathogenesis within *C. elegans* because bacterial toxins produced during infection cause paralysis and death in infected worms [17,18]. We adapted this system to evaluate bacteriophage therapy for multidrug-resistant bacterial infections within the context of a native microbiome. To establish the model, we engineered PA14 to express the orange fluorescent protein mKO_k_ and confirmed expression by fluorescence microscopy. MKO_k_-PA14 cultures exhibited orange fluorescence, whereas wild-type PA14 did not (Figure 1A). Following feeding with mKO_k_-PA14, fluorescence was observed in the intestinal tract of live worms, confirming gut colonization (Figure 1B).

**Figure 1.**
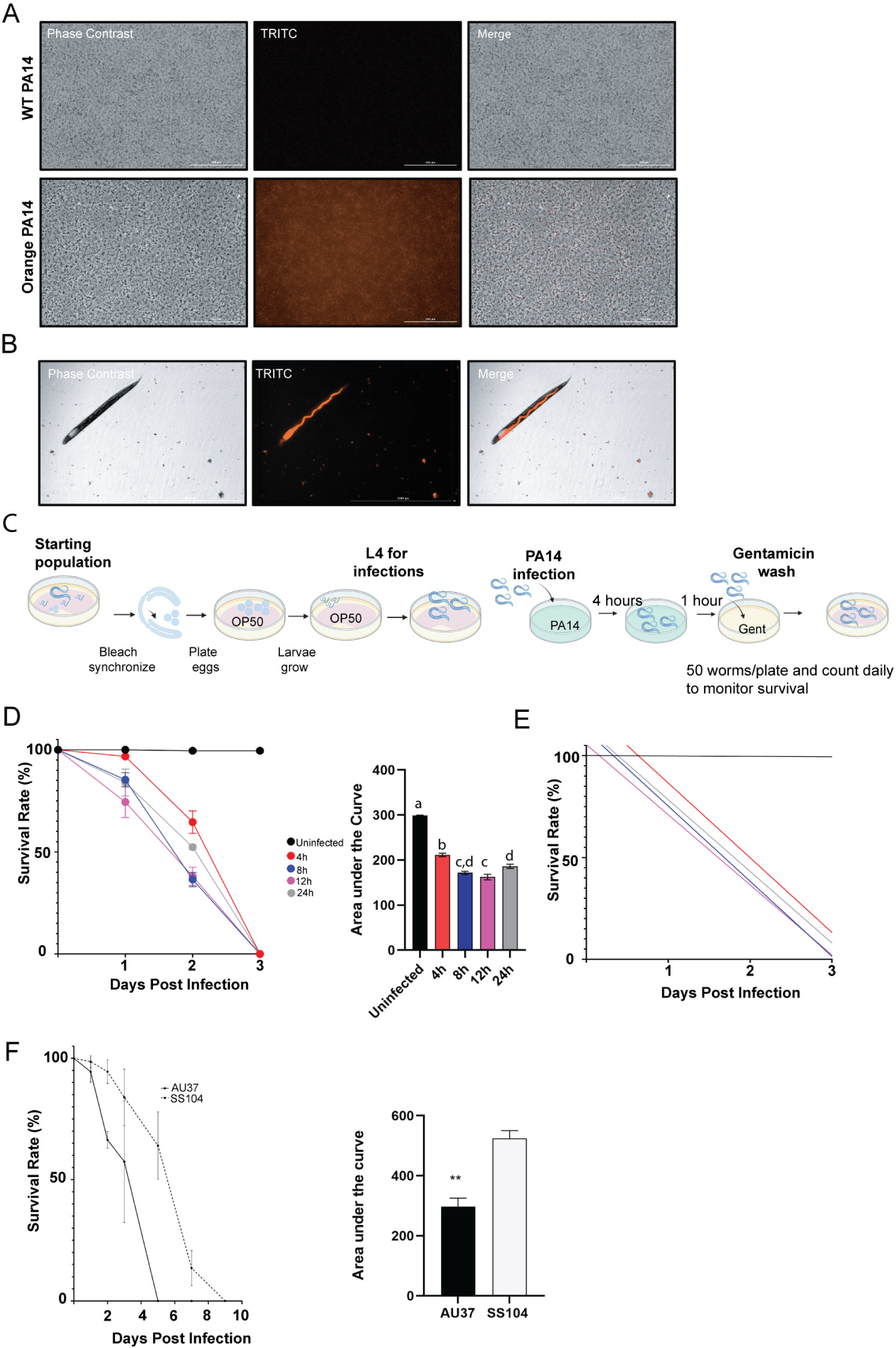
Establishment of a fluorescent PA14 infection model in AU37 *C. elegans*. *Pseudomonas aeruginosa* strain PA14 was engineered to express the orange fluorescent protein mKO_k_ (**A**). The gut of *C. elegans* is colonized with mKO_k_ expressing PA14 following feeding on agar plates (**B**). Schematic overview of the PA14 infection protocol of *C. elegans* (**C**). AU37 worms were exposed to mKO_k_-PA14 for the designated length of time before transfer to OP50 lawns for recovery and daily assessment of survival by microscopy and tactile stimulation. The area under the curve was calculated and used as a measure of the length of survival. There was a statistically significant difference between the area under the curve for 4-hour exposure and all other times (F (4, 10) = 172.3, p < 0.0001, one-way ANOVA with Tukey’s multiple comparisons test) (**D**). A linear regression was performed to model the survival curves observed following various length exposures to PA14 infection (**E**). Temperature restricted reproduction strains AU37 and SS104 were infected with mKO_k_ -PA14 for 4 hours, then 50 worms were transferred to OP50 plates and survival was monitored daily by microscopy. Area under the curve analysis was used to enumerate survival data. AU37 worms exhibited significantly less survival than SS104 worms (*t* = 5.921, *p* = 0.0042, Welch’s *t* test) (**F**).

We next optimized PA14 infection length to achieve gut colonization without causing immediate host mortality during the phage treatment window. Bleach-synchronized worms were fed on OP50 lawns until reaching the L4 stage, and then fed on PA14 for 4, 8, 12, or 24 hours before transfer to OP50 recovery plates for daily survival monitoring (Figure 1C). While all exposure durations resulted in complete mortality, the rate of death was significantly different between treatment groups (Figure 1D, E). Quantification of the area under the survival curve (AUC) revealed a significant reduction in survival for exposures longer than 4 hours (*F* (4,10) = 172.3, *p* < 0.0001, one-way ANOVA with Tukey’s multiple comparisons test) (Figure 1D). Linear regression modeling the slope of the survival curve lines showed that the 4-hour exposure produced a survival slope most like the uninfected control, indicating the greatest therapeutic window for phage intervention (Figure 1E).

Because *C. elegans* strains differ in susceptibility to infections [22], we compared two temperature-sensitive sterile strains to identify a suitable host for PA14 infections and phage treatment studies. Worms were maintained at 25°C to prevent reproduction and ensure uniform infection exposure. Following a 4-hour PA14 challenge, AU37 worms [glp-4(bn2); sek-1(km4) X], which are immunocompromised, showed reduced survival compared to SS104 worms [glp-4(bn2) I] (Figure 1F). AU37 worms exhibited a significantly lower AUC than SS104 worms (296.9 ± 28.3 vs. 524.2 ± 26.0, mean AUC± SE; *t* = 5.921, *p* = 0.0042, Welch’s *t*-test). Therefore, the AU37 strain was selected for use in the model because treatment effects could be detected within a one-week timeframe.

### Lytic phage vB_PaeP_P1G targets PA14 and replicates in the gut of *C. elegans* during infection

Bacteriophages active against PA14 were isolated from wastewater samples collected from treatment facilities in Beacon and Cornwall, NY. Enrichment cultures from three independent samples produced plaques on PA14 lawns but not on *E. coli* OP50 controls, indicating PA14 activity (Figure 2A). Phage activity was further confirmed by monitoring PA14 growth by OD600 growth assay over 48 hours. All three phages significantly reduced bacterial growth relative to untreated controls, as by measured area under the curve (AUC) (1735.7 ± 43.2, 1956.4 ± 91.3, and 1692.3 ± 42.0 vs. 3487.8 ± 46.2; mean AUC ± SE; one-way ANOVA with Dunnett’s multiple comparisons, *F* (3,44) = 206.9, *p* < 0.0001) (Figure 2B, C). We next calculated the bacteriophage virulence using a previously described method [19]. Although virulence scores did not significantly differ among the phages (0.502 ± 0.059, 0.439 ± 0.091, and 0.512 ± 0.042; mean ± SD; *F* (2,32) = 0.7113, *p* = 0.4986), distinct growth kinetics were observed (Figure 2B, C). PA14 treated with vB_PaeP_P1G exhibited rapid growth inhibition followed by later recovery, with a first critical point occurring approximately 170 min after inoculation (Figure 2D). In comparison, vB_PaeS_PAJD-1 P1G and Quinobequin-P09 reached their first critical points at approximately 470 and 610 min, respectively (Figure 2D).

**Figure 2.**
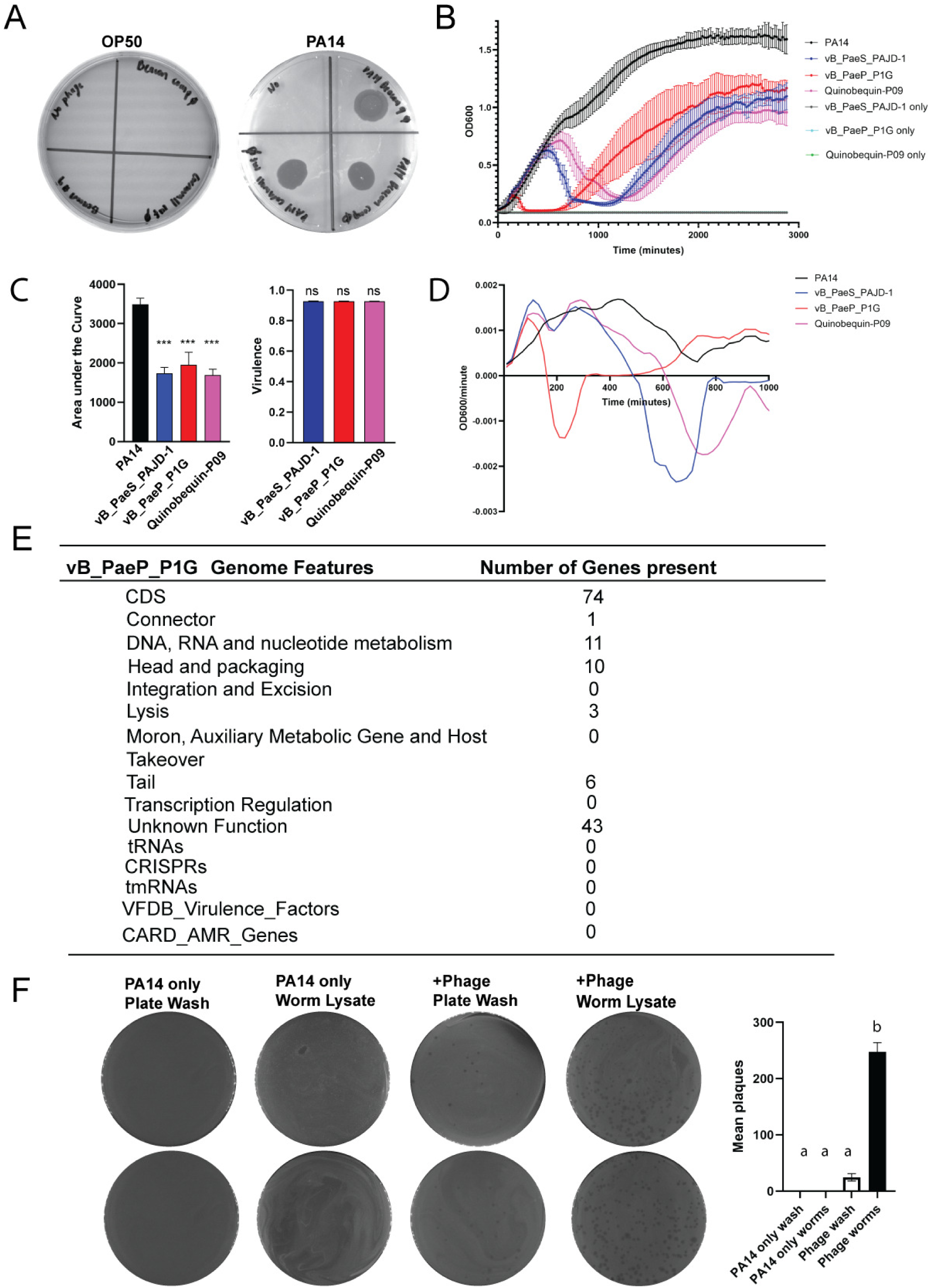
Integration of bacteriophage into the *C. elegans* infection and phage treatment model. Following isolation, purification and expansion of bacteriophages identified in wastewater samples taken from Beacon and Cornwall, NY, 3 phages showed activity against PA14 by plaque assay (**A**) and OD600 growth assay (**B**). The area under the OD600 growth curves was enumerated and used to calculate the virulence for each of the phages compared to the growth of the untreated *PA14* control (black) (**C**). The first derivative was calculated for the OD600 growth per minute to determine the critical points when culture growth changed from positive to negative as a measure of time for phage impact on PA14 growth (**D**). DNA was extracted from phage samples and whole genome sequencing was performed. BLAST searches of the DNA sequences identified the following phages: Pseudomonas phage vB_PaeS_PAJD-1 (94.94% identity, 98% query coverage), Pseudomonas phage vB_PaeP_P1G, 96.71% identity, 100% query coverage), and Pseudomonas phage Quinobequin-P09, 94.63% identity, 97% query coverage. The genome features for vB_PaeP_P1G are shown (**E**). AU37 worms were infected with PA14 and transferred to NGM plates containing OP50 lawns with vB_PaeP_P1G. After 6 days, worms were washed from the plates, lysed by bead beating, and used for incubation with PA14 for plaque assay to quantify plaque forming units (PFU) in the worm gut. Significantly more plaques were observed in worm lysates fed on OP50 plates + vB_PaeP_P1G than in the wash buffer taken from the same plates (*F* (3,4) = 183.8, *p* < 0.0001); One-way ANOVA, Tukey’s multiple (**F**).

Whole-genome sequencing identified the phage isolates as *Pseudomonas* phages vB_PaeS_PAJD-1 P1G (94.9% identity, 98% query coverage), vB_PaeP_P1G (96.7% identity, 100% query coverage), and Quinobequin-P09 (94.6% identity, 97% query coverage). Genomic analysis of vB_PaeP_P1G revealed three lysis genes and no virulence genes, host takeover, or antimicrobial resistance genes, consistent with a lytic lifestyle (Figure 2E). Plaque morphology supported this classification (Fig S1). The remaining phages also contained lysis genes and formed small plaques with zones of bacterial clearing (Table S1). Based on the rapid growth inhibition and lytic nature, vB_PaeP_P1G was selected for therapeutic studies and propagated to a titer of approximately 6.2 × 10^10^ PFU/mL (Fig S2, S3).

To evaluate phage delivery *in vivo*, bleach-synchronized *C. elegans* were infected with PA14 for 4 hours and then transferred to OP50 recovery lawns with or without vB_PaeP_P1G. Six days later, phage abundance was quantified in worm lysates and corresponding wash buffers by plaque assay. No plaques were observed in samples from control worms. In contrast, worms exposed to vB_PaeP_P1G produced more plaques from worm lysate (247.50 ± 23.33; mean ± SE) than the wash buffer from the plate (24.50 ± 6.50) (*F* (3,4) = 183.8, *p* < 0.0001), indicating that the phage was consumed by the worms and replicated within the intestinal tract (Figure 2F).

### Treatment with vB_PaeP_P1G prolongs *C. elegans* death from *PA14* infection but does not rescue

After establishing the infection and treatment model, we next evaluated whether vB_PaeP_P1G improved survival following mKO_k_-PA14 infection. AUC analysis showed that phage-treated worms survived significantly longer than untreated worms, but survival for both groups remained lower than the uninfected OP50 controls (*F* (3,8) = 499.5, *p* < 0.0001; one-way ANOVA with Tukey’s multiple comparisons test) (Figure 3A). Fluorescence microscopy was used to quantify mKO_k_-PA14 intestinal fluorescence in live worms to measure bacterial burden during treatment during another infection. Phage-treated worms exhibited visibly reduced gut fluorescence compared with untreated infected controls at 5 days post infection (Figure 3B). Quantification of the TRITC signal confirmed a significant reduction in mean intestinal fluorescence intensity following vB_PaeP_P1G treatment (*t* = 12.09, *df* = 19.69, *p* < 0.0001, Welch’s *t*-test) (Figure 3C). Together, this demonstrates that vB_PaeP_P1G treatment reduces PA14 colonization of the *C. elegans* gut and improves host survival, but it does not rescue the worms from death (Figure 3B, C).

**Figure 3.**
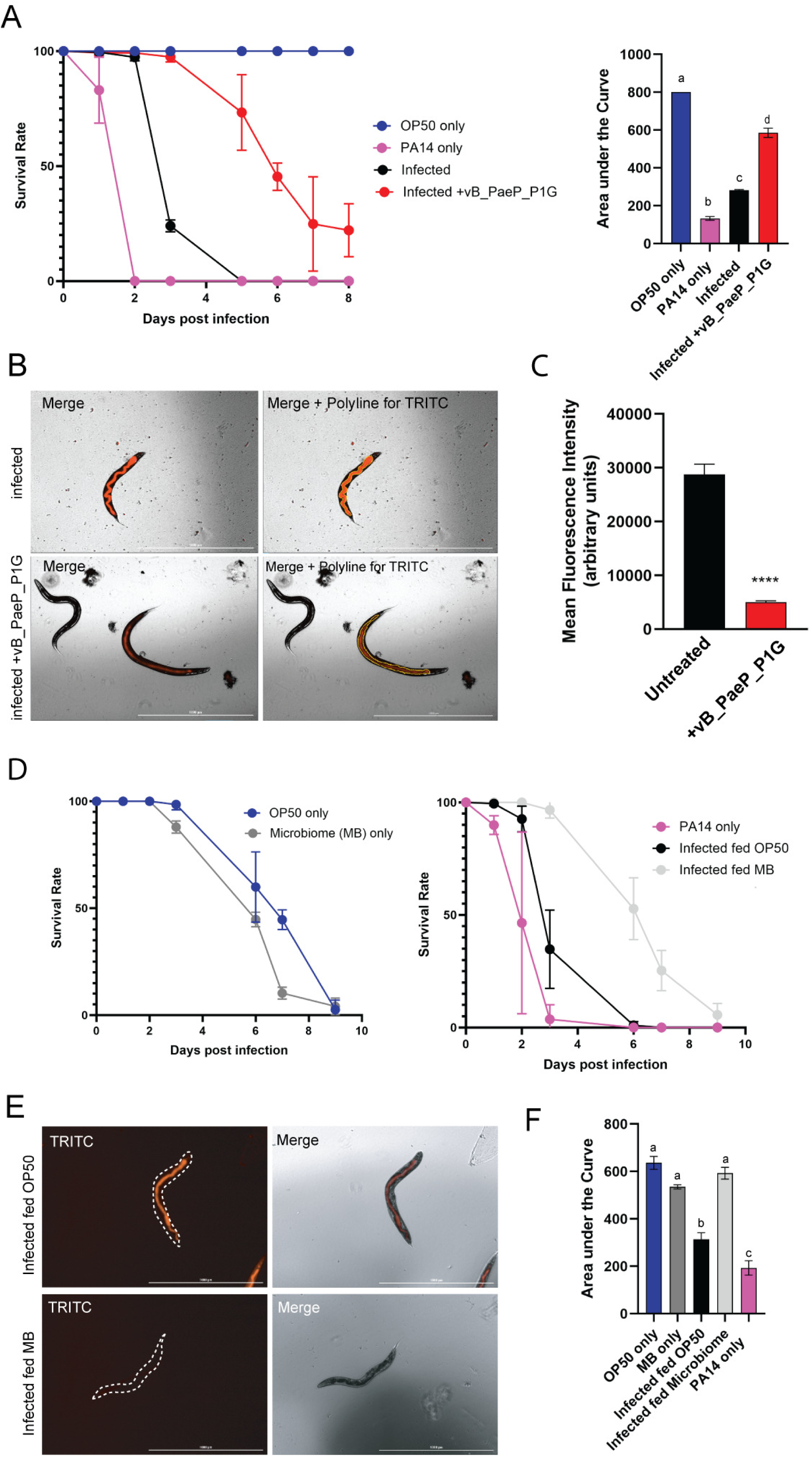
Establishment of the native *C. elegans* microbiome confers protection against PA14 infection. AU37 worms were bleach synchronized and fed on plates containing OP50 until reaching the L4 life stage. Worms were infected with PA14 and transferred to recovery plates containing either OP50 only (infected) or OP50 + vB_PaeP_P1G (infected + vB_PaeP_P1G. Worm survival was monitored daily by microscopy and tactile stimulation. The area under the curve was calculated for the resulting survival curves showing phage treatment improves *C. elegans* survival from PA14 infection but does not rescue them from death (**A**). AU37 worms were infected using the same protocol described above. 5 days post infection, worms were washed from the plates and imaged on the BioTek Cytation V to quantify mKO_k_-PA14 fluorescence in the gut. Representative images of the mKO_k_ fluorescence (TRITC) are shown for untreated infected worms and infected + vB_PaeP-P1G treated worms (**B**). The mean fluorescence intensity of the TRITC channel representing the PA14 infection was calculated showing a reduction in PA14 infection level in the vB_PaeP-P1G treated worms (*n* = 60) at day 5 post infection compared to the untreated control (*n* = 20 worms) (**C**). The difference in sample size between the 2 groups is attributed to the difference in survival from infection at this time point. AU37 worms were bleach synchronized and fed on plates containing the 11 strain *C. elegans* microbiome until reaching the L4 life stage. Worms were infected with PA14 and transferred to recovery plates containing either OP50 only, or the 11-strain microbiome with and without phage. Worm survival was monitored daily by microscopy and tactile stimulation. (**D**). Worms were washed from the plates at day 7 post infection and imaged using the TRITC channel to visualize mKO_k_-PA14 infection. PA14 infected worms fed OP50 are highly infected on day 7, while *PA14* infected worms maintained on CeMbio lawns have limited mKO_k_ fluorescence within the boundary of the worm (dotted white line) (**E**). The area under the curve was calculated for the survival curves showing that the presence of a gut microbiome reduces PA14 infection compared to a single strain alone (infected fed OP50) (**F**).

### The presence of a Microbiome alone reduces PA14 infection burden in *C. elegans*

The microbiome of wild *C. elegans* (CeMbio) has been characterized and is purchasable from the Caenorhabditis Genetics Center (CGC) [16]. We introduced microbiomes to the infection model by feeding worms members of the CeMbio community prior to PA14 infection and phage treatment. The identity of 11 of the 12 CeMbio strains purchased from the CGC was confirmed by PCR and then used to establish microbiome lawns for worm colonization (Fig S4). The presence of a gut microbiome during infection significantly improved survival from PA14 in both a 3-strain pilot study and the complete 11-strain microbiome (Figure 3D, Fig S5; *F* (4,10) = 58.25, *p* < 0.0001, one-way ANOVA with Tukey’s multiple comparisons test). Infected worms fed OP50 exhibited significantly greater mKO_k_-PA14 fluorescence at day 7 than worms colonized with the CeMbio collection, indicating less infection when the microbiome is present (Figure 3E). Survival analysis further demonstrated that PA14-infected worms fed OP50 had significantly lower AUC values (313.7 ± 28.0; mean ± SE) than OP50-only controls (636.4 ± 27.2), microbiome-only controls (535.0 ± 8.6), or infected worms maintained on the microbiome (592.6 ± 25.1) (Figure 3F). Although infected worms colonized with the microbiome ultimately died, their survival was comparable to uninfected control groups throughout most of the experiment.

We observed reduced survival in the uninfected microbiome control group after day 4. Because several CeMbio strains have been reported as opportunistic pathogens, we tested each strain individually in AU37 worms. *Chryseobacterium scophthalmum* (JUb44) and *Ochrobactrum vermis* (MYb71) caused the greatest reduction in survival (Fig S6), suggesting that opportunistic pathogens within the microbiome contribute to the decline in survival observed in uninfected control groups (Figure 3D).

### Bacteriophage treatment does not significantly impact the composition or diversity of the microbiome

Bacteriophages target specific host strains; however, removal of one species from a microbial community can perturb the overall community structure from shifts in available resources or changes in interspecies competition [13]. If key taxa are removed, dysbiosis can result, decreasing the microbial diversity and leaving vacancy for pathogens to invade [12, 13]. We show that vB_PaeP_P1G is not active against CeMbio strains by agarose overlay and OD600 growth assays (Fig S7). While vB_PaeP_P1G does not kill CeMbio strains, it is unclear how phage treatment may impact the microbiome community within the worm when used to combat infection. AU37 worms colonized with the CeMbio community were infected with mKOk-PA14 and treated with or without vB_PaeP_P1G. After 3 days, bacterial DNA was isolated from worms and analyzed by 16S rRNA sequencing (Figure 4A). Comparison of the starting inoculum and plate-grown lawns revealed community shifts during growth, most notably a reduction in *Pseudomonas olverans* B and increases in *Acinetobacter guillouiae* (MYb10) and *Sphingobacterium paramultivorum* (BIGb0170), indicating differential fitness among community members under lawn growth conditions (Figure 4B, Table S2).

**Figure 4.**
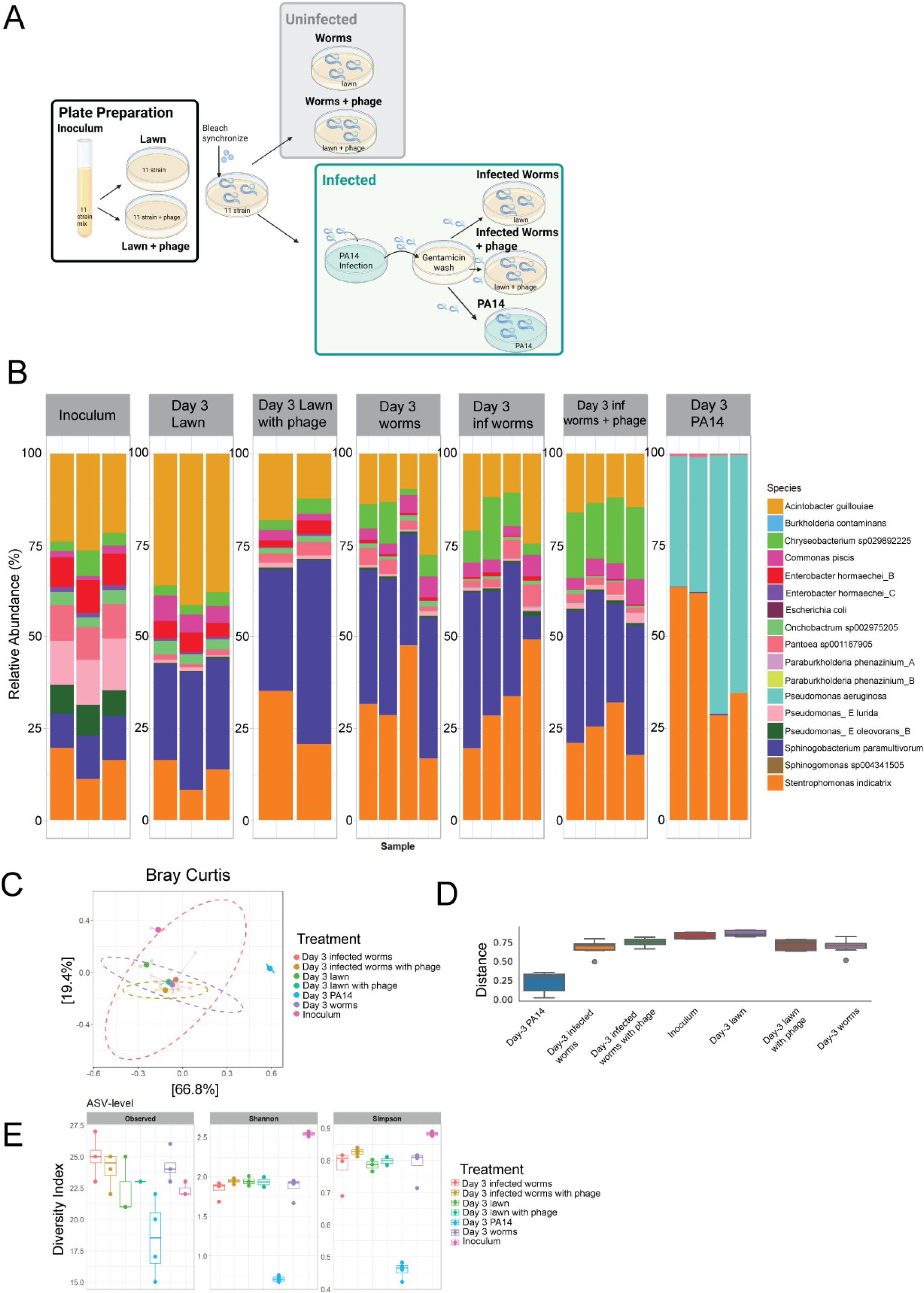
Phage treatment does not significantly alter the *C. elegans* gut microbiome composition during PA14 infection. Schematic showing how the DNA samples were generated during the PA14 infection process for 16S sequencing of the microbiome. AU37 worms were bleach synchronized and fed on plates containing the 11-strain *C. elegans* microbiome until reaching the L4 life stage. Worms were infected with PA14 and transferred to recovery plates containing the microbiome with and without phage. After 3 days, worms were washed from the plates, DNA was extracted, and the full length 16S rRNA gene was sequenced with PacBio sequencing (**A**). Microbial composition plots showing the mean percent relative abundance for each of the members of the *C. elegans* microbiome are shown (*n* = 2-4 biological replicates per condition) (**B**). Bray Curtis analysis comparing the beta diversity in each of the samples revealed that the most spatially separated sample by principal component analysis is the Day 3 PA14 (blue). All other samples cluster near one another showing similarity (**C**). Enumeration of the distances between samples when plotting the principal component analysis generated by Bray Curtis (**D**). Comparison of amplicon sequencing variants (ASVs) by Observed, Simpson, and Shannon as measures of the alpha species diversity indices further supports that the inoculum and PA14 only samples are the most different samples and that the others show no significant difference in microbiome composition (**E**).

The intestinal microbiome of the worms was dominated by *Sphingobacterium paramultivorum* (BIGb0170), *Stenotrophomonas indicatrix* (JUb19), and *Acinetobacter guillouiae* (MYb10), with similar relative abundances across infected, phage-treated, and uninfected worms (Figure 4B). Bray–Curtis beta diversity analysis demonstrated substantial overlap among microbiome communities regardless of infection or phage treatment status (Figure 4C, D). The greatest divergence was observed in worms maintained on PA14 for 3 days without recovery, whose microbiomes were dominated by *Pseudomonas* and *Stenotrophomonas* and exhibited reduced representation of other taxa (Figure 4B).

Alpha diversity analyses further supported the stability of the microbiome during phage treatment. As expected, the inoculum mixture exhibited the greatest diversity, whereas worms continuously maintained on PA14 displayed the lowest richness and diversity (Figure 4E, Table S3). In contrast, neither lawn nor worm samples showed notable differences in Shannon diversity between phage-treated and untreated groups. While no differences in diversity were detected, the mean relative abundance of PA14 in day 3 infected worms with phage sample was lower than in the untreated day 3 infected worms (Fig S8). Pairwise-PERMANOVA identified no significant differences between treatment groups (Table S4). Together, these results indicate that vB_PaeP_P1G selectively reduces PA14 while preserving the overall composition and diversity of the native *C. elegans* microbiome.

### Infection level during bacteriophage treatment does alter the native microbiome of *C. elegans*

Having established that vB_PaeP_P1G targets PA14 without altering the microbiome at day 3, we next examined whether infection level influences the microbiome composition. AU37 worms colonized with the 11-strain CeMbio community were infected and treated as previously described, and DNA samples were collected at day 3 (low infection exhibiting minimal mortality) and day 6 (high infection, approximately 50% mortality) for 16S sequencing (Figure 3D). The experimental microbiome community remained stable once it transitioned from the inoculum to the experimental environment. While the inoculum was enriched for *Pseudomonas lurida* (MYb11) and *Pantoea nemavictus* (BIGb0393), growth on agar selected for communities dominated by *Sphingobacterium paramultivorum* (BIGb0170), *Stenotrophomonas indicatrix* (JUb19), and *Acinetobacter guillouiae* (MYb10). This pattern was maintained in both lawn and worm samples (Figure 5A). Worms maintained on PA14 alone exhibited markedly reduced community complexity and were dominated by *P. aeruginosa* and *S. indicatrix* across all time points, with PA14 comprising 70–93% of the community (Figure 5A, Table S5). Day 0 worm samples, collected immediately after the 4-hour infection period, revealed strong initial selection for PA14. Over time, *S. indicatrix* increased in abundance, indicating that it was the most persistent CeMbio member during competition with PA14.

**Figure 5.**
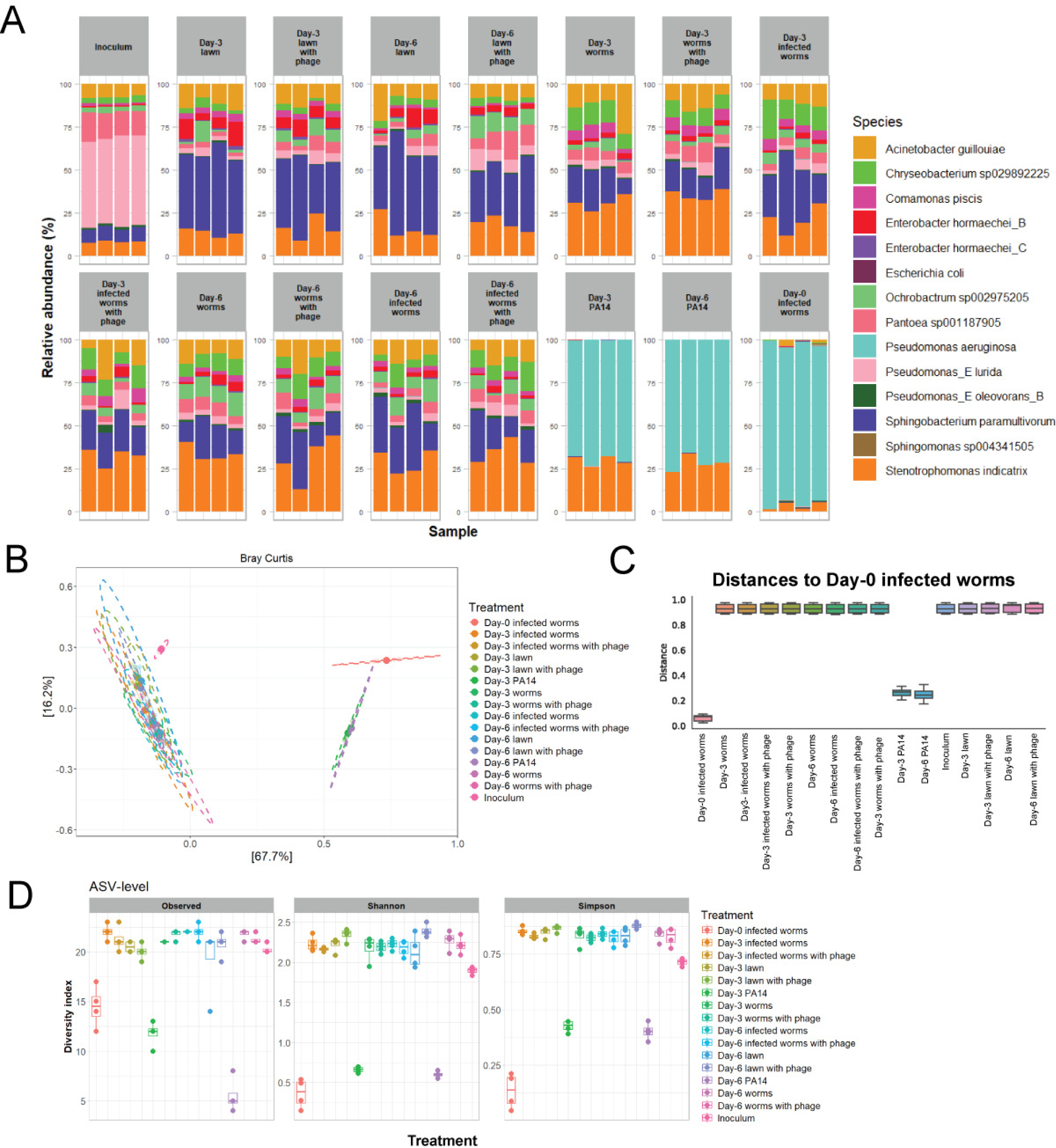
The microbial communities are similar in composition despite treatment status and infection level. AU37 worms were bleach synchronized and fed on plates containing the CeMbio collection until reaching the L4 life stage. Worms were infected with PA14 and transferred to recovery plates containing the microbiome with and without phage. After 3 days (low infection) and 6 days (high infection), lawns and worms were washed from the plates, DNA was extracted, and the full length 16S rRNA gene was sequenced with PacBio sequencing. Microbial composition plots showing the mean percent relative abundance for each of the members of the *C. elegans* microbiome are shown (*n* = 4 biological replicates per condition) (**A**). Bray Curtis analysis comparing the beta diversity in each of the samples revealed that the most spatially separated samples by principal component analysis are the Day 0 worms and the PA14 samples at Day 3, and Day 6. All other samples cluster near one another showing similarity, with the spheres indicating the 95% confidence interval overlapping on top of one another (**B**). Enumeration of the distances between points on the principal component analysis generated by Bray Curtis analysis. The Day 0 worms are most like the Day 3 and Day 6 PA14 samples, as these all are closest together. All other samples are roughly equivalent distance from the Day 0 worms and cluster just under 1. (**C**). The alpha diversity index represented by plotting the amplicon sequencing variants (ASVs) by Observed, Shannon, and Simpson diversity further supports that the Day 0 worms and Day 3 and Day 6 PA14 samples are the most similar. Once the infection has been established, the microbiome diversity stabilizes and remains unchanged despite infection or treatment status (**D**).

Bray–Curtis analysis and alpha diversity metrics confirmed that PA14-only worms contained the least diverse microbial communities and clustered separately from all other samples (Figure 5B–D). Microbiome-colonized worms and lawn samples clustered together regardless of infection status, treatment, or sampling day (Figure 5B, C; Fig S9). The alpha diversity remained similar among all microbiome-containing samples, while the PA14-only worms exhibited the lowest richness and diversity (Figure 5D, Table S6). Pairwise PERMANOVA detected no significant differences between groups (Table S7). The phage-treated worms exhibited lower relative abundances of PA14 than untreated worms at both day 3 and day 6 (Fig S10). Treatment with vB_PaeP_P1G reduces PA14 burden across multiple infection levels and preserves the microbiome composition and diversity, indicating that phage treatment does not induce dysbiosis within this model.

## Discussion

In this study, we hypothesized that the specificity of phages will prevent bacteriophage treatment from disrupting the balance of microorganisms comprising the microbiome, providing a potential benefit to phage-based treatments over antibiotic usage. Phage therapy relies upon identification or synthesis of phages targeting the pathogen of interest. While finding or designing phages may be challenging for some infections, phage therapy provides an alternative treatment for multidrug resistant infections where existing antibiotics are ineffective. Currently phage therapies have been employed for compassionate use to treat infection with ESKAPE pathogens [11]. Specifically, phage cocktails have successfully been used to treat *P. aeruginosa* infections in burn wounds, chronic endocarditis, chronic otitis media and chronic lung infections [11]. A low dose 12 phage cocktail used in the PhagoBurn study reduced the bacterial burden and produced less adverse effects than the standard of care control group [20]. While the phage cocktail reduced the bacterial burden more slowly than the gold standard treatment, the overall reduction was greater and highlighted the promise for phage therapies with additional optimization [20].

Successful clinical experiences, along with stalled antibiotic discovery in the time of accelerated resistance, have led to revisiting phage therapy as a treatment option for MDR infections. Before phage therapy can be employed as a widespread practice, a thorough understanding of how phage administration impacts the microbiome is required. Here we establish a new model system integrating the host, pathogen, phage and microbiome within the same organism to address this question. We show that treatment of PA14 infection with bacteriophage targeting PA14 does not alter the microbiome of *C. elegans*. This is the first example of bacteriophage therapeutic treatments being integrated into a model organism along with a complete microbiome and establishes a pipeline to test phage-pathogen interactions for other ESKAPE pathogens.

Decreased microbial diversity has been reported in patients with hospital acquired infections such as *C. difficile* and sepsis [21]. There is limited understanding to date about how phage treatment within these patient populations may impact microbial diversity. Previous investigations on how phages impact the microbiome have been carried out using a gnotobiotic mouse model [14, 15]. Like the *C. elegans* model, the gut microbiome of gnotobiotic mice is sterile and can be reconstituted with organisms of interest. In these studies, the microbiome was colonized with representative members of human gut microbiome and then virus-like particles (VLP) purified from human fecal samples targeting organisms within the microbiome were administered to represent a “phage attack” [14, 15]. Within this model there are changes in the microbiome following phage treatment, but the underlying differences in the experimental setup compared to those herein may explain the opposing results. The identity of the VLPs administered to the gnotobiotic mice is not known, but target species found in the human microbiome, so removing bacteria by phage treatment created a new ecological niche to be filled by resident bacteria. In the *C. elegans* model, we have introduced a pathogen to colonize alongside the microbiome and then administered phage treatment targeting the pathogen and then assessed changes following removal of the pathogen (Figure 2A, Fig S6). This difference in experimental design has improved our understanding of what may happen during therapeutic phage treatment.

Using the *C. elegans* model fed on the CeMbio diet, we observed that the dominant species of the gut microbiome are *Sphingobacterium paramultivorum* (BIGb0170), *Stentropomonas indicatrix* (JUb19), *Acintobacter guillouiae* (MYb10), regardless of phage treatment status or infection level (Figure 4B, Figure 5A), which is consistent with the reported literature [16, 22]. Additionally, we observe fitness consequences for some strains of the *C. elegans* microbiome (JUb44), which aligns with identified opportunistic pathogens within the CeMbio collection [23, 24].

While we developed a new model to recapitulate therapeutic phage treatment within the context of a complete native microbiome, this is not the first use of *C. elegans* to assess phage-bacteria interactions. A liquid culture model was previously established to deliver both prophylactic and therapeutic phage treatments to *C. elegans* to improve worm survival from *Escherichia coli* 131, *E.coli* 311, *Klebsiella pneumoniae* 235, and *Enterobacter cloacae* 140 infections [25]. Like our observations, administration of bacteriophages in liquid culture improves worm survival from infection but does not rescue the entire population from death (Figure 3A) [25]. The liquid culture model provided greater improvements to *C. elegans* survival rates (>90% survival for prophylaxis) compared to our plate administration model, but because the infection is established when phage is also present in the media, it difficult to assess if the lower infection is due to phage action inside the worms or in the liquid culture itself before infection is established [25]. For this reason, we opted to use the plate administration model to deliver the therapeutic phage dose, despite the previous success in prolonging worm survival with the liquid culture model. Furthermore, phage treatment was effective in reducing infection with *Burkholderia pseudomallei* and *Salmonella enteritidis* in the *C. elegans* model when administering phages on the bacterial lawns for consumption [26, 27]. Here, we integrate the microbiome into this model of infection and treatment for the first time. Like previous studies, we show that simply introducing a microbiome to the experimental model of infection in *C. elegans* results in decreased pathogen burden (Figure 3D, E) [28, 29].

One limitation of our experimental model is that we are using *pmk-1* deficient worms (AU37) to explore the impact of infection and phage treatment. The PMK-1/p38 MAPK signaling pathway is the primary innate immune response in *C. elegans*, making these worms highly susceptible to infections. While it has been shown that disruption in this signaling pathway promotes higher bacterial burdens both for pathogens and the microbiome establishment within AU37 worms, it does not impact the alpha and beta diversity [30]. Within our infection model with fluorescent PA14, we observe total death in the population of AU37 worms by 3 days, while immunocompetent SS104 worms survive infection up to 10 days (Figure 1F). This timeline to death for our mKO_k_-PA14 strain is like survival rates reported elsewhere [31, 32]. Treatment with vB_PaeP_P1G prolongs worm survival but does not rescue the population from death, possibly because a single phage type was used for treatment. It has been well established that multiple phages capable of targeting the same bacterial strain are ideal for construction of cocktail treatments, and efforts are currently underway for continued phage isolation and characterization [11]. While the complex microbiome of higher organisms contains bacteria, viruses and fungi; however, this work provides a starting hypothesis that phage treatment may not impact microbiomes in other systems. Furthermore, we provide a model that can accelerate the development of targeted bacteriophage therapies against MDRs with limited effect on microbiomes.

## Materials and Methods

### PA14 Culture and Engineering Fluorescence

mKO_k_ expressing PA14 was engineered using a *Pseudomonas* specific allelic exchange protocol with plasmids gifted from Dr. Carey Nadell (Text S1 for detailed method) [33] . Screening for fluorescent PA14 colonies following sucrose counterselection, was performed on dilutions of overnight cultures using the Biotek Cytation 5 Cell Imaging Multimode Reader (Agilent, Santa Clara, CA USA) with phase contrast and TRITC filter cubes. The images were processed and deconvolved using the Gen5 Software (RRID:SCR_017317).

### *C. elegans* Imaging on Cytation V

*C. elegans* (AU37) were fed on NGM plates with mKO_k_ -PA14 lawns for 4 hours before transfer to recovery NGM plates with OP50 lawns for 5 days. Worms were washed from the plates with M9 buffer at the desired timepoint for imaging and transferred to a 24 well plate at various dilutions so they were spatially separated for imaging. Worms were imaged with the 4x objective for phase contrast and the TRITC filter cube for fluorescence of mKO_k_ –PA14. The images were processed and deconvolved using the software available in the Gen5 Software.

### Bacteriophage Isolation and Propagation

Raw sewage samples collected from wastewater treatment facilities in Beacon and Cornwall, NY, were used for screening bacteriophage activity against PA14 by double agar overlay and OD600 growth assay as described in [6]. Areas of clearance on PA14 lawns by agarose overlay were isolated and phages were propagated by infection of PA14 cultures grown shaking at 250 RPM (at OD600 = 0.1) at 37°C until the bacteria were fully lysed. OD600 was checked every hour until it fell below 0.05. Lysates were cleared by centrifugation (10 min at 3400 x g) and filter sterilized with a 0.22 μm filter to remove residual bacteria. Final phage titers (in Plaque forming units (PFU)/ mL) were determined by counting plaques using the double agar overlay method.

### Phage DNA Isolation and Sequencing

The 3 samples containing bacteriophage with activity against PA14 were propagated in PA14 as described above. 5 mL of each sample (all containing PFU > 10^8^ PFU/mL) was concentrated by centrifugation using Amicon Ultra Centrifugal Filter Units (MillporeSigma, Burlington, MA, USA) with 100K kD pore size. Following centrifugation, the filter unit was washed twice with 1 mL of DNAse/RNAse Free water. To remove residual bacterial DNA and RNA, the samples were treated with 1 μL of DNase I (1 U/ μL) (RNase Free DNase Set cat #79254, Qiagen, Germantown, MD, USA) and 1 μL of DNase and Protease free RNase A (10 mg/mL) (cat# EN0531, ThermoFisher Scientific, Waltham, MA, USA) and incubated at 37°C for 90 minutes [34]. The DNase and RNase were inactivated by adding 20 mM ETDA (20 μL of 0.5 M stock). The samples were then treated with 1.25 μL Proteinase K (cat# 1014023 Qiagen,Germantown, MD, USA) and incubated for 1.5 h at 56°C to digest the phage capsid protein [34]. Following this, a Phage DNA Isolation Kit (Norgen BioTek Corp, Thorold, ON, CA) was used to extract DNA following the kit instructions. The elution step was performed using DNase/RNAse free water. 21 μl (minimum concentration 39 ng/ μL) of DNA was sent to CD Genomics (Shirley, NY, USA) for Phage Whole Genome Sequencing and Assembly according to the company instructions for sample submission.

### Quantification of mKO_k_ – PA14 fluorescence in *C. elegans*

*C. elegans* were infected with PA14 and transferred to plates with OP50 lawns only or OP50 with vB_PaeP_P1G for recovery. 5 days post infection, worms were washed from plates with M9 buffer as previously described and transferred to a 24-well microplate to dilute the worm solution, so the bodies were spatially separated. The BioTek Cytation V Cell Imaging Multimode Reader was used to capture images for mKO_k_ fluorescence quantification. Worms were imaged with both Phase Contrast and TRITC (556, 600) using the 4X objective. All worms in the well were imaged. The Cellular Analysis Tool found in “Process/Analyze features” was used to perform image quantification. The channel for Cellular Analysis was set to TRITC, and the cell analysis parameters, such as Threshold, Minimum Size, and Maximum Size were altered such that the bacterial fluorescence was captured in the worms, but not in the surrounding external environment. The “Polyline Tool” was used to outline the worms to reduce background signal from mKO_k_ washed from the infected plates. The mean fluorescence intensity for every worm in the captured images was determined by the Gen5 program and exported to Excel.

### Sequencing Analysis Pipeline for PacBio Sequencing of 16S rRNA genes

Raw sequencing data (FASTQ files) obtained from SeqCenter were used to perform a complete QIIME2 16S analysis. Two different QIIME2 workflows were utilized simultaneously: a traditional QIIME2 workflow, detailed below, and a NextFlow HiFi Full Length 16S analysis.

The traditional QIIME2 workflow was as follows: import, demultiplex, denoise, feature tables, assigning taxonomy, phylogenetic analysis, diversity analysis, and differential abundance. The FASTQ files were imported and used to create a QIIME2 artifact. Using QIIME2 demux, the sequences were demultiplexed, grouped together by their individual barcodes. Next, the sequences were denoised with QIIME data2, producing high quality amplicon sequence variants (ASVs) by removing sequencing errors, noise, and chimeras from the raw sequences. Feature tables were prepared and summarized using QIIME feature-table summary to identify whether samples were required to exclude or set a rarefaction depth. Taxonomic assignment was performed using QIIME feature-classifier. A new classifier was developed and trained to best suit the sequence data used for this pipeline using the QIIME feature-classifier extract-reads. The new classifier was then used to sort all the ASVs into their taxonomy and then downstream phylogenetic, diversity, and differential abundance analyses were possible. Using QIIME phylogeny alight-to-tree-mafft-fasttree, unrooted and rooted phylogenetic trees were produced. Alpha and beta diversity analyses were performed using QIIME diversity core-metrics-phylogenetic to generate plots for alpha diversity (Shannon’s Diversity, Observed Features, Faith’s Phylogenetic diversity) and beta diversity plots (Bray-Curtis, Jaccard Distance, and unweighted UniFrac).

However, using the trained classifier and the default QIIME2 classifier failed to provide species level resolution. For this reason, the NextFlow HiFi Full-length 16S analysis was utilized. Unlike the traditional QIIME2, this specific NextFlow workflow utilized two classifiers simultaneously: QIIME2 Naïve-Bayes classifier and the VSearch classifier. Unlike Naïve Bayes, VSearch is better suited for high quality full-length PacBio sequences. The NextFlow workflow was downloaded and run using Lawrence Livermore National Lab’s Mammoth cluster. The outputs of the NextFlow workflow provided species level resolution for all taxa.

RStudio and Vega Editor generated plots for taxa barplot, Bray-Curtis, Alpha Diversity, and ANCOM-BC using the QIIME2 files produced by NextFlow analysis. in RStudio, the following QIIME2 analyses were imported into phyloseq class: dada-ccs_table, rooted phylogenetic tree, taxonomy, and the metadata. ggplot and the phyloseq object were then used to create the figures.

### Quantification and Statistical Analyses

GraphPad Prism (RRID: SCR_002798) was used for all statistical analyses. A standard t test was used when measuring the difference between two experimental groups. One-way ANOVA with Tukey’s multiple comparisons test was used when more than three groups.

## Additional methods

See Text S1 in the supplemental material for detailed methods describing the PA14 allelic exchange protocol, bacteriophage plaque assay, nematode growth media (NGM) plate preparation, *C. elegans* maintenance and bleach synchronization, PA14 infection of *C. elegans*, the generation of *C. elegans* survival curves, and PCR confirmation of CeMbio strains.

## Acknowledgements

This work was supported by funding from the Defense Threat Reduction Agency through a grant awarded to Dr. Andrew Kick (R.0005103.242.12) and salary support from Dr. F. John Burpo. We thank Dr. Carey Nadell for the generous sharing of the plasmid and the technical advice required to make mKOk expressing PA14. We also would like to thank Dr. Caroline Amoroso for technical advice and instruction on how to establish the *C. elegans* model. Phage DNA sequencing and downstream genome assembly was a paid service carried out by CD Genomics (Shirley, NY, USA). PacBio sequencing of the 16S rRNA genes for the microbiome study was a paid service performed by SeqCenter (Pittsburgh, PA, USA). The Caenorhabditis Genetics Center was instrumental in maintaining the stocks for purchase of all worm and bacteria strains needed for this research. BioRender was used to design schematic illustrations for experimental design.

